# Phylogenomics analyses of all species of Swordtails (Genus *Xiphophorus*) highlights hybridization precedes speciation

**DOI:** 10.1101/2023.12.30.573732

**Authors:** Kang Du, Yuan Lu, Mateo Garcia-Olazabal, Ronald B. Walter, Wesley C. Warren, Tristram Dodge, Molly Schumer, Hyun Park, Axel Meyer, Manfred Schartl

## Abstract

Hybridization has been recognized as an important driving force for evolution, however studies of the genetic consequence and its cause are still lagging behind in vertebrates due to the lack of appropriate experimental systems. Fish of the central American genus *Xiphophorus* were proposed to have evolved with multiple ancient and ongoing hybridization events, and served as a valuable research model in evolutionary biology and in biomedical research on human disease for more than a century. Here, we provide the complete genome resource and its annotation of all 26 *Xiphophorus* species. On this dataset we resolved the so far conflicting phylogeny. Through comparative genomic analyses we investigated the molecular evolution of genes related to melanoma, for a main sexually selected trait and for the genetic control of puberty timing, which are predicted to be involved in pre-and postzygotic isolation and thus to influence the probability of interspecific hybridization in *Xiphophorus*. We demonstrate dramatic size-variation of some gene families across species, despite the reticulate evolution and short divergence time. Finally, we clarify the hybridization history in the genus *Xiphophorus* genus, settle the long dispute on the hybridization origin of two Southern swordtails, highlight hybridizations precedes speciation, and reveal the distribution of hybridization ancestry remaining in the fused genome.

## Introduction

The debate in evolutionary biology on the role and importance of hybridization and postzygotic hybrid sterility genes in the process of speciation remains unsettled. For decades, hybridization was thought to be rare in animals and perhaps particularly so in vertebrates^1^. However, this view was overturned by an increasing number of genomic studies over the last two decades. With this growing appreciation for the ubiquity of hybridization, researchers have become interested in the evolutionary consequences of hybridization, particularly for adaptation and as a mechanism of speciation^1–5^. While examples for reticulate evolution^6,7^ are increasing, evidence for hybrid speciation in vertebrates is still rare.

*Xiphophorus* is a genus of Central American freshwater fishes^8^. To date, 26 species have been described that occur in various freshwater habitats in the Atlantic drainages of Mesoamerica, from Northern Mexico to Guatemala. Historically, they have been classically divided into four groups according to their geographic distribution: the Northern and Southern platyfishes and the Northern and Southern swordtails. Their striking morphological variation and the possibility to experimentally produce inter-species hybrids^9,10^ render *Xiphophorus* an ideal vertebrate model system to address questions related to the role of hybridization in phenotypic evolution and speciation. In previous work, for two species an origin from a hybridization event was proposed ^11–13^ and a transcriptome based survey revealed evidence for reticulate evolution^14^. Moreover, there are several known ancient^15^ and contemporary hybrid zones in this group^16,17^.

Although hybridization, especially within *Xiphophorus*, appears to be more frequent than previously thought, the evolutionary impact of hybridization will be shaped by the extent to which hybrids between lineages survive and reproduce and thereby introduce novel genetic material into different lineages. In nature both pre- and postzygotic isolation mechanisms impact the formation and persistence of hybrids. Prezygotic isolation can be mediated by species-specific differences in courtship and mating behavior, among other mechanisms^18,19^. *Xiphophorus* fish have been intensively studied in part because of their dramatic sexually selected ornaments^20–22^. One of the most well studied sexually selected traits in swordtails – first noted by Darwin -- is a colorful extension of the ventral rays of the caudal fin referred to as the “sword”^23^.This trait is found in all species of Southern swordtails and most Northern swordtails, and absent in platyfishes^8^. The sword ornament is used in courtship displays. It is highly attractive to females in many species^24,25^ and serves as an important reproductive barrier in species that have lost the sword^26^. A “preexisting bias” hypothesis postulates that the female preference is ancestral and facilitated the evolution of the trait. Knowledge about a mono- or polyphyletic origin of the sword in combination with a precise species tree is required to validate this hypothesis^27^. Candidate “sword development genes” responsible for an overgrowth of fin rays have been proposed for two loci on chromosome 13^16,28^, but their true involvement in production of this spectacular trait remains unproven.

Differences in life history traits involved with reproduction may be barriers to hybridization. In several species of *Xiphophorus* the onset of male sexual maturity is genetically determined by so-called puberty loci (*P*) on the sex chromosomes^29^. In two species it was shown the *P* loci harbors various wildtype and defective mutant alleles encoding melanocortin 4 receptors, Mc4r^30^, a gene that is known to be involved in metabolic regulation, obesity and onset of puberty in mammals^31^. Although a molecular mechanism for the action of dominant negative versions of Mc4r in regulating the onset of the reproductive period and associated traits such as adult size, courtship and dominance behaviors, has been revealed^32–34^, the phylogenetic distribution of this system and its evolution are not known.

The evolutionary outcomes of hybridization also depend crucially on genetic interactions that occur when the genomes of two divergent lineages are combined. Postzygotic isolation in *Xiphophorus* became a textbook example for a “speciation gene”^35–37^, or a genetic interaction that generates reproductive barriers between lineages. This emerged from studies on spontaneous development of melanoma among select *Xiphophorus* interspecies hybrids^38,39^. Several laboratory hybrid crosses, as well as natural hybrid populations of *Xiphophorus* species^40^, develop malignant pigment cell lesions and serve as biomedical models for understanding the genetic etiology of the human disease in translational studies^41^. A mutant copy of an epidermal growth factor receptor *(egfrb*) allele, *xmrk*, has been detected in *X*. *maculatus*^42,43^ and acts as a melanoma-inducing oncogene. Normally, in *X*. *maculatus* the *xmrk* oncogene is kept in check by an epistatic trans-acting regulatory locus *R/Diff*, while in hybrids, crossing conditioned-elimination of *R/Diff* leading to *xmrk*, dysregulation causes cancer. Previous studies indicated that *xmrk* is absent in some *Xiphophorus* species and that not all interspecies crosses result in melanoma-bearing hybrids^44,45^. The evolutionary relevance of the *R/Diff-xmrk* oncogene/tumor suppressor gene situation in some but not all species, and questions about a monophyletic vs convergent evolution of this trait are currently unknown.

To formally test evolutionary hypotheses for the occurrence and transmission of these and many other important traits in *Xiphophorus* species we require comprehensive and high-quality genomic resources and an accurate phylogeny. Reference genomes for several representative species of the genus have been generated, but for most of the species annotated genome assemblies are missing. Several phylogenies based on mitochondrial sequences and/or partial nuclear genomic sequences have been forwarded^12–14,27^ and led to conflicting results with respect to species placements and ancient gene flow, or hybrid origins of the species. We sequenced, assembled and annotated the genomes of all known *Xiphophorus* species as well as three undescribed taxa of the genus *Xiphophorus* for which genomes had not been determined previously. From the most complete genomic dataset for this genus we generated the mitochondrial and nuclear phylogenies of all *Xiphophorus* species and studied the evolution of the genomes and selected genes. We find evidence of extensive hybridization during the evolution of *Xiphophorus* based in both current and ancestral lineages.

## Results

### Genome assembly and annotation

From the genus *Xiphophorus* five chromosome-level assembled genomes (*X*. *maculatus*, *X*. *hellerii* and *X*. *couchianus*, *X*. *birchmanni*, *X*. *malinche*) have been published^40,46^. Here, we add twenty-nine additional genomes to these existing genomic resources, which were sequenced using Illumina, 10X, PacBio and/or Hi-C techniques in this study (Supplementary Table S1). This now provides a complete genome resource of all *Xiphophorus* species, including all 26 previously identified species (https://www.ncbi.nlm.nih.gov/Taxonomy/Browser/wwwtax.cgi?id=8082), three undescribed taxa, *X*. *sp* I, *X*. *sp* II, *X*. *sp* III, two new strains of *X*. *maculatus*, and a reference genome for the species *Priapella lacandonae* as outgroup.

The resulting assemblies ranged from ∼636Mb to 720Mb (close to the estimated genome size^47^), with a scaffold N50 ranging from ∼2Mb to 32Mb. BUSCO analyses demonstrate that each assembly covers ∼92-97% of the 4584 well-conserved Actinopterygii genes (Supplementary Table S2). Using RepeatModeler and RepeatMasker^48^, we found ∼30-35% of the content of the *Xiphophorus* assemblies are made up of transposable elements (TEs) (Supplementary Table S5). When aligning each assembly to *X*. *maculatus* (GCA_002775205.2), the coverage is ∼84-92% (Supplementary Table S3). Average divergence between sequences within the *Xiphophorus* genus were all lower than ∼2.5% (Supplementary Table S4). However, as expected from both hybridization and variation in constraint and the strength of background selection^49,50^, sequence divergence varies along the genome. Each chromosome has regions of higher and lower sequence conservation. Alternating regions of high or low divergence on a chromosome are observed in a consistent pattern across the whole genus (Supplementary Fig. S1, S2).

Using a custom annotation pipeline^51^, we annotated ∼22-25k protein coding genes (PCGs) in each assembly, improving the BUSCO assessment of completeness by ∼1% (Supplementary Table S6). Based on the similarity of protein sequences, these PCGs were clustered into 26,982 gene families (Supplementary Fig. S3). Out of these gene families, we find evidence that 353 families underwent a significant change in the size of the family during evolution of the genus *Xiphophorus* . These include olfactory receptors, odorant receptors, retinol dehydrogenase 12 and melanocortin 4 receptors (*mc4r*) (Supplementary Fig. S4; Supplementary Table S7).

Those genomes, which are assembled at chromosome level (*X*. *birchmanni*, *X*. *malinche*, *X*. *couchianus*, *X*. *maculatus*, *X*. *hellerii* and *P*. *lacandonae*), show in general a high conservation of synteny with the exception of the two platyfish species, *X*. *maculatus* and *X*. *couchianus*. They have several translocations and inversions compared to the karyotypes of the Northern and Southern swordtails (Supplementary Fig. S5).

### Gene evolution

The availability of the genomes from all swordtails and platyfishes allowed us to determine the species specific presence or absence and origin of genes that are of importance for *Xiphophorus* as a model for speciation and human diseases.

#### *xmrk* melanoma oncogene

The *xmrk* gene was originally found in the Southern platyfish, *X*. *maculatus*. From the whole genome information of all species, we were able to retrieve *xmrk* sequences in a total of nine platyfish and Northern swordtail species, but not in any of the Southern swordtails. Due to the high degree of conservation of *xmrk* and *egfrb* (its proto-oncogenic precursor) coding sequences, several nodes of the gene tree are only poorly supported (Supplementary Fig. S6). Despite this, a monophyletic origin of *xmrk* is visible.

#### Sword candidate genes

The sword candidate gene *kcnh8*^28^ is present in *Xiphophorus*, as in other non-polyploid teleosts as a single copy gene. The gene tree largely follows the species tree and no amino-acid changes were detected that correlated with absence or presence of a sword. The same situation was found for another sword candidate gene *sp8* postulated in an independent study^16^.

#### Defective copies of *mc4r*

The onset of reproductive maturity in males of two species (*X*. *nigrensis*, and *X*. *multilineatus*) is regulated by the number of mutant *mc4r* copies clustered on the Y chromosome (historically referred to as the *P*-locus^30^). These copies are defective for transmitting the intracellular signal from *mc4r* due to changes in the carboxy terminal amino acid sequence. From the genome assemblies we were able to identify multiple mutant copies, which are predicted to be defective, also in *X*. *maculatus*, *X*. *xiphidium*, *X*. *birchmanni*, *X*. *malinche*, *X*. *evelynae* and *X*. *sp* III, but not in any of the Southern swordtails. This may suggest an origin of the *P*-locus system at the base of the Northern swordtail and platyfish clades, similar to the situation of the gene duplication that produced *xmrk*. Further studies are required to reconstruct the evolutionary history of the different mutant alleles and the variation in copy numbers between and within species.

### Discordance of mitochondrial and nuclear phylogenetics

Discordance in phylogenetic trees constructed from partial mitochondrial and nuclear genomic data in *Xiphophorus* has been previously reported in several studies ^12,14,27^ and is not uncommon in many taxa^52–54^. Such discordance can indicate a history of gene flow or can be generated by processes such as incomplete lineage sorting. With our complete genome resources, including both mitochondrial and nuclear nuclei, we are now able to revisit this question at a genome-wide scale. For the mitochondrial phylogenetic tree, sequences of all 37 mitochondrial genes were aligned across the species independently, and then concatenated for a maximum-likelihood estimation in RAxML. The tree was also reconstructed using Bayesian approaches as an alternative method, which generated a nearly identical topology (Supplementary Fig. S7). For the nuclear sequences, we reconstructed a phylogenomic tree based on a ∼342Mb long whole genome alignment (WGA) using stringent criteria (see Material and Methods) using RAxML under the GTR+Gamma model. All nodes of the tree were fully supported as indicated by the 1,000-bootstrap test. According to mitochondrial and nuclear phylogenomics the genus *Xiphophorus* is divided into three monophyletic groups: platyfishes (unifying the previously distinguished Northern and Southern platyfishes), Northern swordtails and Southern swordtails. However, the platyfishes were grouped with Southern swordtails in the mitochondrial tree but with Northern swordtails according to the nuclear phylogeny (Fig. 1). This incongruence was also visible in trees in two previous studies, but not further analyzed^12,14,27^.

**Figure 1.**
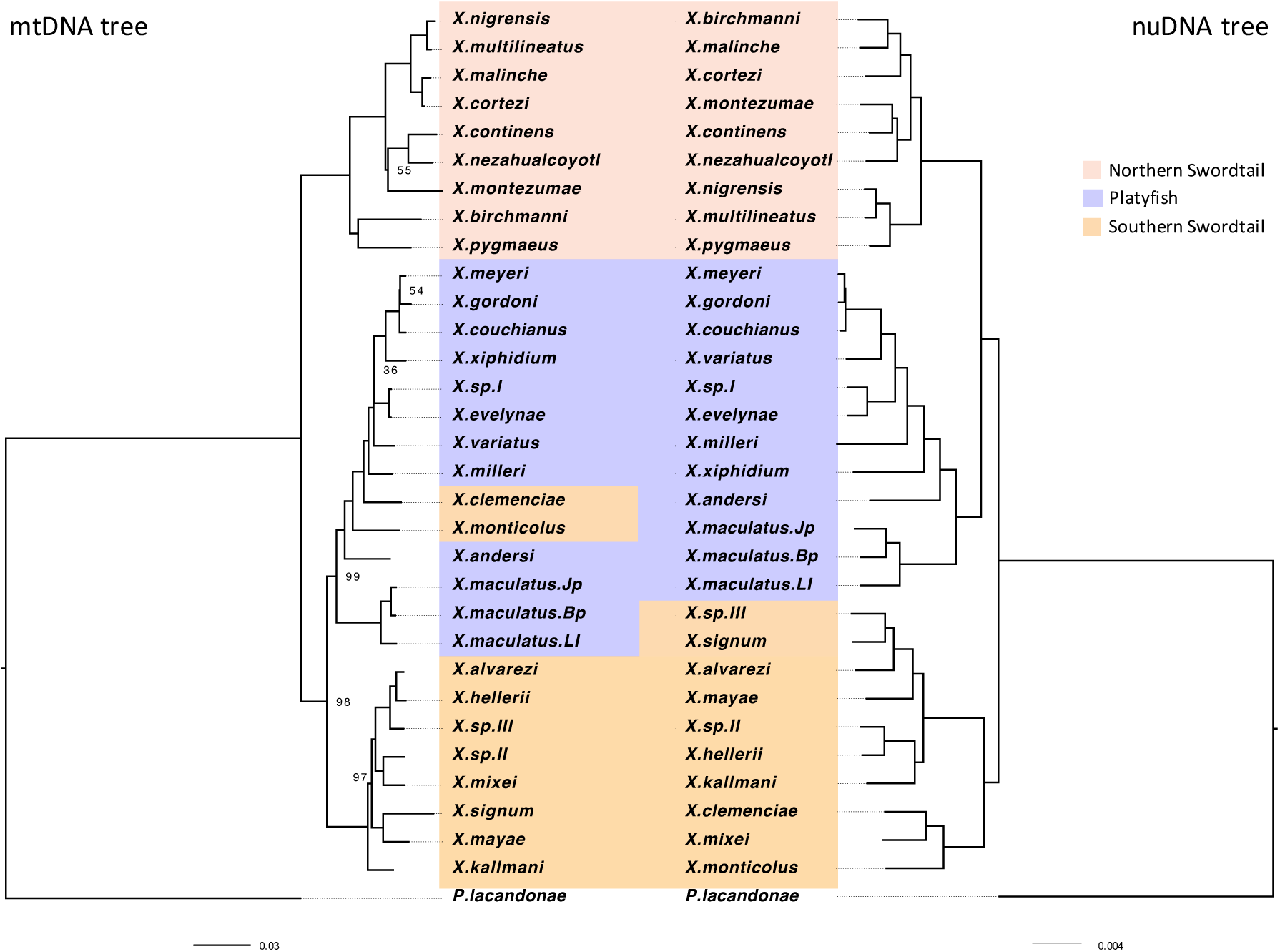
Phylogenetic trees constructed using maximum-likelihood method for the mitochondrial genome (left) and nuclear genome sequences (right). Numbers on the nodes represent bootstrap support values; nodes without number means that they are 100% supported.

Our result also confirmed previous reports of the conflicting placement of two southern swordtail species, *X*. *clemenciae* and *X*. *monticolus*. As expected from morphological traits, in the nuclear tree they are placed in the southern swordtail clade, but in the mitochondrial tree they are nested within the platyfish-lineage.

Based on the protein coding region for reconstructing the nuclear phylogeny, we built maximum-likelihood trees using 3,259 one to one orthologous genes with their concatenated protein sequences, coding sequences and fourfold degenerate sites (4DTV). The resulting phylogenetic estimates were almost identical to the WGA tree (Supplementary Fig. S8), except *X*. *xiphidium* and *X*. *alvarezi* were placed in slightly different positions within their respective clade.

Discordance between gene trees and species trees can be generated by the coalescent process or by historical gene flow. We investigated this further using coalescenct methods to build a nuclear phylogeny. To restrict sequence sampling to nonrecombinant loci, we randomly sampled 19,111 WGA blocks that are 20 kb away from each other and selected a one-kb window from each block to build the individual trees (Supplementary Fig. S9A). After removing trees with poor supporting value (see Materials and Methods), we built a coalescent phylogeny using ASTRAL-III. Notably, in this analysis, *X*. *clemenciae*, *X*. *monticolus* and *X*. *mixei* were placed outside of the Southern swordtails and basal to the Northern swordtails and platyfishes as a separate clade (Supplementary Fig. S9B).

Overall, the phylogenetic tree of the whole genus *Xiphophorus*, and the usage of information from whole genome sequencing, are largely in accordance with placements of the species from previous morphology-based and partial sequence-based trees, except for *X*. *continens*. Two studies found a close association of *X*. *continens* with *X*. *pygmaeus*^13,14^, while all our trees place *X*. *continens* as sister taxon to *X*. *montezumae*. Our result is consistent with another recent study on Northern swordtails^55^. Our whole genome re-sequencing of the material used in the conflicting studies revealed mistyping of the material (data not shown). Our analysis also strengthens the position of the Northern platyfishes species as being a crown group within the Southern platyfishes. Consequently, a distinction between both groups is no longer justified and we refer to the whole clade as “platyfishes”.

### Extensive reticulate hybridization during the evolution of the genus *Xiphophorus*

The presence of “platyfish-mitochondria” associated sequences in Southern swordtail species implies that two species, *X*. *clemenciae* and *X*. *monticolus*, have experienced hybridization with the platyfish lineage during their evolution. This is in accordance with earlier data from partial mitochondrial sequences and some nuclear loci^12,27^.

Phylogenetic discordance can be generated by hybridization, but also by technical factors such as base compositional biases or rate differences, or the coalescent process via incomplete lineage sorting (ILS). It is possible to distinguish the effect of hybridization from ILS by implementing *f_4_*-ratio statistics using the program Dsuite with the “*f*-branch” method and WGA data^56^. This revealed the nuclear genome of the ancestor of *X*. *clemenciae* and *X*. *monticolus* contained more than 10% admixture proportions derived from the ancestral lineage of platyfishes and Northern swordtails (Fig. 2), consistent with a hybrid origin of *X*. *clemenciae* and *X*. *monticolus*. Results also show that this hybridization must have occurred before the split of *X*. *clemenciae*, *X*. *mixei* and *X*. *monticolus*. A possible hybrid origin of *X*. *mixei* was not previously considered since mitochondrial haplotypes from these species are not discordant with the species tree. A further hybridization event was also detected between *X*. *mixei* and *X*. *monticolus* (Fig. 2). The considerable amount of sequence originating from outside of the Southern swordtails clade may explain the more basal position of the *X*. *clemenciae*/*X*. *monticulus*/*X*. *mixei* subclade in the coalescent tree (Supplementary Fig. S9B).

**Figure 2.**
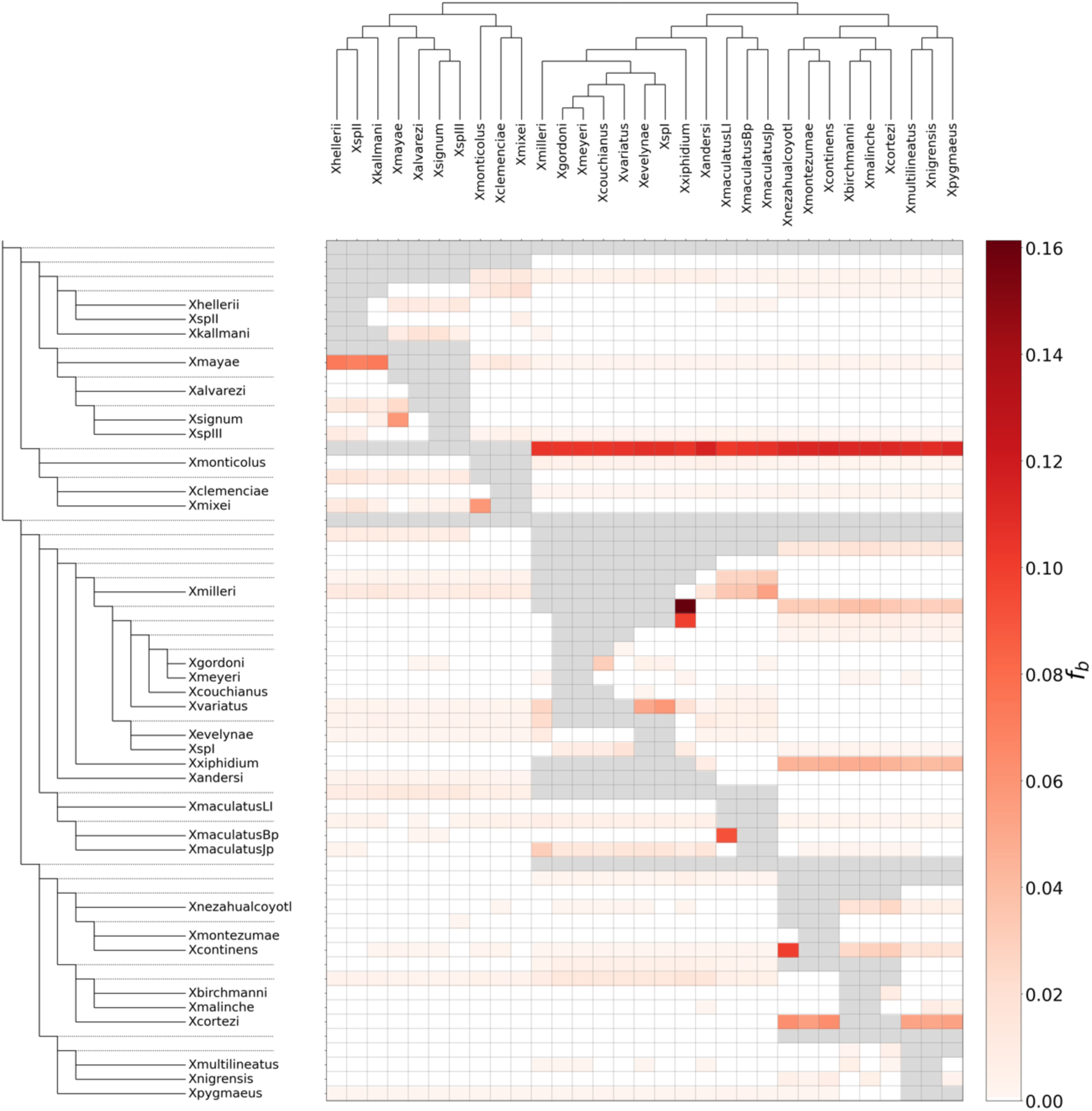
Heatplot showing the estimated admixture fraction between two species on the x and y axis. Species are listed on the x and y axis in the species tree. The values in the matrix (*f*_b_) were calculated using Dsuite with the *f*-branch statistic, referring to excess allele sharing between the branch identified on the expanded tree on the y axis (relative to its sister branch) and the species identified on the x axis, indicating the hybrid origin of these alleles.

Apart from the hybrid ancestry of the *X*. *clemenciae*, *X*. *mixei* and *X*. *monticolus* subclade our analyses uncovered several other hybridization events in the genus *Xiphophorus*. A strong signal was detected in the ancestral branch leading to *X*. *gordoni*, *X*. *meyeri*, *X*. *couchianus*, *X*. *variatus*, *X*. *evelynae* and *X*. *sp* I with *X*. *xiphidium*. Other admixture events are detected between the *X*. *maculatus* Bp and LI populations, and between *X*. *signum* and *X*. *mayae*. In agreement with a recent phylogenomic study focusing on the Northern swordtails^55^, we observe gene flow between *X*. *cortezi* and several other Northern swordtail species and between *X*. *continens* and *X*. *nezahualcoyotl* . Finally, we detect substantial gene flow between *X*. *mayae* and *X*. *kallmani*, *X*. *hellerii* and *X*. *sp* II.

### Admixture signals across chromosomes

We used Dsuite to explore the distribution of introgressed loci across chromosomes for species with hybridization history. As control, we also included *X*. *alvarezi* (Fig. 3A) which did not show any signals of hybridization (Fig. 2). Results show different distributions of hybridization-derived regions on different chromosomes for different hybridization events (Fig. 3). For *X*. *signum* (Fig. 3I), hybridization-derived regions are clustered in the middle of Chromosome 2, 15 and 18, while for other species, these regions are spread over the entire genome (Fig. 3). Conspicuously, a common admixture pattern is shared within the clade of *X*. *clemenciae*, *X*. *monticolus* and *X*. *mixei* (Fig. 3B-D) and among *X*. *couchianus*, *X*. *variatus*, *X*. *evelynae* and *X*. *sp* I (Fig. 3E-H), respectively. Local differences were found when the pattern of *X*. *couchianus* and *X*. *variatus* (Fig. 3EF) was compared to that of *X*. *evelynae* and *X*. *sp* I (Fig. 3GH), potentially reflecting the influence of the secondary hybridization with *X*. *xiphidium* (Fig. 2).

**Figure 3.**
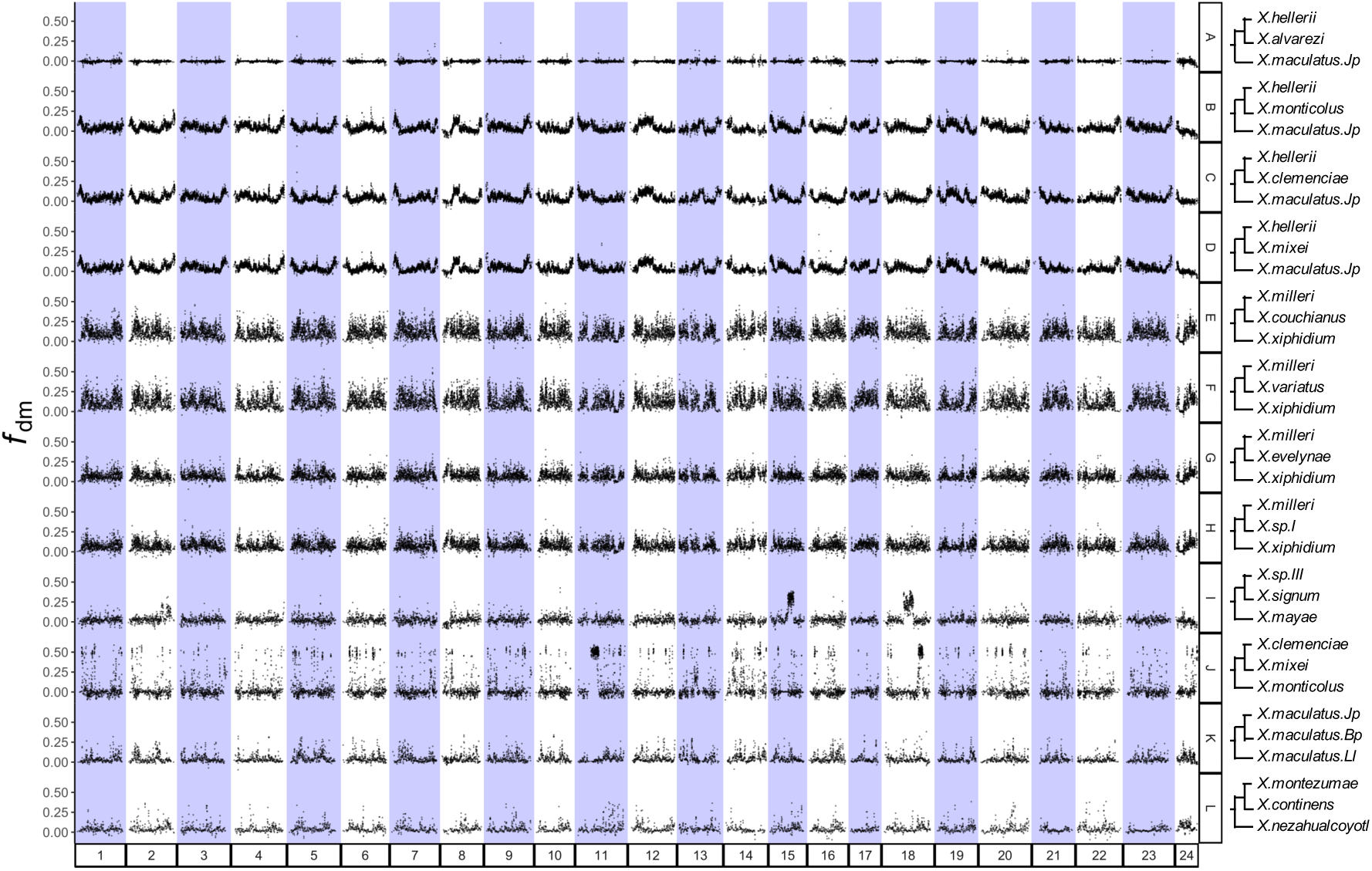
Dot plots revealing genome regions derived from hybridization for species with or without hybrid history. Trees on the right are in the format of ((P1,P2),P3) where admixture signal between P2 and P3 was investigated. Values (*f*_dM_) on the y-axis were calculated using Dsuite with modified *f*_d_ statistics developed by Malinsky et al. (2015), referring to the fraction of excess alleles in slide-windows sharing between P2 and P3 than between P1 and P3, indicating hybridization origin of the genome region. Statistics were averaged over windows of 100 informative SNPs, moving forward by 50 SNPs. Row A refers to a genome without hybridization resources.

Hybridization-derived regions shared by multiple species provide an opportunity to investigate the evolutionary fate of those regions following hybridization. We identified those regions in *X*. *clemenciae* using AU-test with WGA data. Among all the WGA regions that passed AU-test (AU *P*-value >0.95, see Methods and Materials), 49.2% (95% CI: 49.0-49.3%) of them yielded the topology placing *X*. *clemenciae* and *X*. *maculatus* together (Topology 2, Supplementary Fig. S10AB), significantly more than those placing *X*. *hellerii* and *X*. *maculatus* together (3.1%, 95% CI: 3.1-3.2%; Topology 3, Supplementary Fig. S10AB). The estimation of the admixed partition is higher than the f4 ratio test, however, it gives qualitative support for *X*. *clemenciae* being a hybrid lineage and also allows us to identify hybridization-derived regions. Those hybridization-derived regions show a lower sequence divergence (k2p value) than the others when comparing the DNA sequence between *X*. *clemenciae* and *X*. *monticolus* (Supplementary Fig. S10C). Using the same method with protein coding gene (PCG) alignments, we also identified significantly more PCGs supporting Topology 2 than that supporting Topology 3 (Supplementary Fig. S10D). Functional enrichment analysis using shinyGO^57^ suggested no function is over-represented in these PCGs. Pairwise sequence comparison between *X*. *clemenciae* and *X*. *monticolus* revealed lower synonymous substitution rates (dS) and higher ratios of nonsynonymous substitution to synonymous substitution (dN/dS) for these PCGs (Supplementary Fig. S10EF).

## Discussion

*Xiphophorus* has been used as a model to study questions from basic processes in evolutionary biology to biomedicine and human disease for over a century. In this study we have sequenced, assembled and annotated genomes for 19 *Xiphophorus* species and two new *X*. *maculatus* strains, thereby generating complete genomic resources for the whole genus *Xiphophorus* for the first time. Together with five earlier reported genomes^16,46^, we provide new insights on micro and macroevolutionary processes within the genus, generate the first whole-genome based phylogeny for all species, characterize the history of hybridization and investigate the patterns of hybridization-derived regions along the genome.

### Gene evolution

Combining our robust species tree with the phylogeny of *xmrk* and *egfrb* supports a single origin of *xmrk* at the base of the Northern swordtail and platyfish clades. However, many species in the Northern swordtail and platyfish clades do not have *xmrk,* suggesting that it has been lost several times independently.

The melanoma inducing action of *xmrk* is controlled by a co-evolved unlinked tumor suppressor locus on chromosome 5 for which several candidate genes have been suggested^40,58,59^. Phylogenomic studies on congruence of allelic differences in those genes with the presence or absence of a sword may help to determine about the most likely gene candidate and to understand its mode of action.

The sword candidate gene *kcnh8* encodes a potassium channel. Potassium channels are found frequently to be mutated in other fish that show overgrowth of fins^60–64^. While in those species the respective gene is a duplicate from the teleost-specific whole genome duplication or a polyploidization event that obviously underwent subfunctionalization for regulating fin growth, we find only one copy of *kcnh8* in all *Xiphophorus* species and in *Priapella*. Because of the pleiotropic function of this gene (e.g., in neuronal cell), a structural change like those observed in other fin mutant channels would compromise also *kcnh8* in other cells. Thus, it may be more likely that a regulatory change is the basis of sword formation.

Another conclusion comes from the complete phylogeny of the genus that confirms the sword has evolved prior to speciation of the swordless platyfish^27^. Such an origin is not compatible with the pre-existing bias hypothesis which postulates that the sword evolved in ancestrally swordless species due to the preference of females for such a male ornament.

Mutant *mc4r* alleles are predicted to have a strong effect on dynamics of growth and development. In *X. nigrensis* and *X. multilineatus*, *mc4r* is a major determinant of the age of puberty and the other characters linked to this trait (e.g. male size, ornamentation, courtship behavior and territory establishment and defense)^32–34^. Breeding experiments have indicated that the timing of the onset of puberty is also heritable in *X*. *variatus*, *X*. *milleri*, *X*. *montezumae* and *X*. *cortezi*^29^. The absence of mutant *mc4r* copies in these genomes may be due to incomplete assemblies. However, a previous study showed that in *X*. *hellerii* the polymorphism in onset of maturity, and the connected adult male size, are not determined by the interaction of wildtype and mutant isoforms^32^. Further, it was shown that large males have much higher hypothalamic expression of *mc4r*, pointing to the same regulatory pathway but a different molecular mechanism underlying the same phenotypic polymorphism.

### Microevolution of *Xiphophorus* genomes at the genus level

Genetic variation and genome evolution in vertebrates were investigated widely either within populations or across distant species, yet genome comparisons between species within a genus that diverged only a few million years ago are few. According to earlier studies^8,44^, *Xiphophorus* species radiated roughly around 5 MYA. Our study now provides complete genomic resources for these closely related species to study genomic variation and evolutionary patterns over short evolutionary timescales.

We aligned all genomes to that of *X*. *maculatus*, which is assembled on chromosome scale based on the most extensive short and long-read data collection including a meiotic map^46,65^. The alignments between chromosome-level assemblies revealed a high degree of conserved synteny. Moreover, in two species of *Xiphophorus* it was found that they have conserved recombination maps^66^. This may indicate that patterns of constraint and background selection may be generally shared across the genomes of these species. Indeed, we identify shared patterns of variation in diversity and divergence between species (Supplementary Fig. S1). Similar distributions of these highly diversified regions were also observed when the genomes were aligned to that of *X*. *hellerii* (Supplementary Fig. S2). When mapping homology across six species with chromosome-level genomes including outgroup *Priapella*, we found inversions and translocations of chromosome segments in platyfishes, despite conserved local synteny over the whole genome having been maintained for at least 25 million years (Supplementary Fig. S5). These bursts of rearrangements in the platyfish clade are striking and may be linked to a higher lineage specific structural turnover of the karyotype. Future research on those rearrangement may aid evaluating the impact of chromosome structure on speciation (see ^67,68^) in the genus *Xiphophorus*.

It is worth noting that the phylogenomic results revealed larger genetic distances between strains of *X. maculatus* than among some species, particularly *X*. *meyeri*, *X*. *gordoni* and *X*. *couchianus* or *X*. *multilineatus*, *X*. *nigrensis* and *X*. *pygmaeus* (Fig. 1). Species in *Xiphophorus* have been usually identified and defined by differences in morphology, physiology, lifestyle and ecological adaptations. Intuitively, with more pronounced phenotypic differences, stronger interspecific genetic variation may be expected. However, our genomic data show a case where there is more genetic variation between populations of *X. maculatus* than between phenotypically divergent species. It will be interesting to investigate whether there is cryptic isolation between these *X. maculatus* populations using crossing experiments and tests for assortative mating. It will also be important to explore whether other factors such as higher mutation rates or larger historical population sizes could have contributed to these patterns. Lineage-specific bursts of trait evolution vs. stasis linked to fast or slow diverging genomic features has been observed and well studied between distant and long diverged lineages^69^. *Xiphophorus* offers the opportunity to investigate this now at a microevolutionary scale.

The size of certain gene families also varies significantly across the genus *Xiphophorus* (Supplementary Table S7). While these patterns have been previously reported for *paox* and *mc4r*^46^, we identify exciting new patterns of gene family expansion in our genome-wide dataset. Perhaps most notable given the importance of sexual selection in *Xiphophorus* is the identification of diversification in gene families associated with vision and olfactory functions. These gene families may allow the divergent adaptations among different species, for instance, in olfactory preferences^70^, which are crucial premating barriers in *Xiphophorus* species^71,72^. The pronounced lineage-specific dynamicsgene family size provides new clues for trait diversification in the genus.

### Phylogenomics and hybridization

Our phylogenomic analyses divided *Xiphophorus* into three monophyletic groups: platyfishes, Northern swordtails and Southern swordtails, in agreement with previous studies on a subset of species^13,14^. With the complete dataset, we were able to resolve the placement of several species that were previously disputed. For example, *X*. *xiphidium* was placed at the base of the platyfish clade in a phylogenetic tree constructed by using eleven nuclear loci^12^. Our results suggest this misplacement may be caused by restricted nuclear data available or previously undetected hybridization event(s) in the species history of *X*. *xiphidium* (Fig. 2). Similarly, gene flow between *X*. *signum* and *X*. *mayae* (Fig. 2), may have contributed *to X. signum* being identified as the sister lineage to *X*. *alvarezi* in previous work^14^. Together with another recent study^55^, we consider mistyping of material for *X*. *continens* in previous studies^13,14^.

Discordance between mitochondrial and nuclear phylogenies can indicate instances of past gene flow. In our phylogeny based on the complete mitochondrial genome the platyfishes were grouped with Southern swordtails, but placed with Northern swordtails in the nuclear tree. Hybridization origin of platyfishes could be an appealing explanation. However, genomic segments shared between platyfishes and Southern swordtails (∼6.5%) are not significantly more than those shared between Northern swordtails and Southern swordtails (∼4.7%), hence ILS cannot be excluded to account for this mito-nuclear discordance.

The mitochondrial genomes of two Southern swordtail species, *X*. *clemenciae* and *X*. *monticolus*, are perfectly nested within the platyfish clade, making us revisit the hypothesis that they may have arisen through hybridization^11–13,27^. Past research proposed that this mito-nuclear discordance could be generated by ILS since whole genome sequences did not support admixture in the nuclear genome^14,73^. Here, using *P. lacandonae* as the outgroup, we demonstrate that the amount of potential hybridization-derived loci of *X*. *clemenciae* is significantly higher in the nuclear genome than expected for ILS alone (Supplementary Fig. S10BD). The discordance with past research may be driven by the fact that this hybridization event appears to be more ancient than previously thought, localizing before the divergence of platyfishes and Northern swordtails(Fig. 2).

Our result also suggests that the hybridization leading to *X*. *clemenciae* and *X*. *monticolus* is the same event, which occurred before the split of the *X*. *clemenciae*, *X*. *monticolus* and *X*. *mixei* clade. Different from the other two species, *X. mixei* has its mitochondrial haplotypes assigned to the Southern swordtails. Multiple independent fish sampling in the field from previous studies have implied that *X. monticolus* and *X. clemenciae* harbor mitochondria of only the “minority” parental species ^11–13^. Thus, the presence of Southern swordtail mitochondria in *X*. *mixei* may be explained by backcrossing of hybrid fish with a female Southern swordtail (Fig. 2).

More generally, complete genome resources of all *Xiphophorus* species give us a broader view of the hybridization history among all current and ancestral lineages. Our pan-genus genomic information uncovered the common ancestral hybridization event in the lineage of *X. clemenciae*/*X. mixei*/*X. monticolus* was contributed by an ancestor of platyfishes and Northern swordtails (Fig. 2), suggesting that the event that led to mitonuclear discordance was much more ancient than previously hypothesized (Supplementary Fig. S11). Our phylogenomic analysis clearly reveals that the hybridization event preceded the formation of the three Southern swordtail species. In addition, we also saw hybridization between *X*. *nezahualcoyotl* and *X*. *continens* (Fig. 2)^55^, which is more prominent than the previously reported hybridization between *X*. *nezahualcoyotl* and *X*. *cortezi*^14,15^, making this lineage of particular interest for unraveling a complex history of hybridization in that group. Finally, our study discovered ancient hybridizations between *X*. *xiphidium* and the ancestor of *X*. *gordoni*, *X*. *meyeri*, *X*. *couchianus*, *X*. *variatus*, *X*. *evelynae* and *X*. *sp* I (Fig. 2), unveiling a profound influence of *X*. *xiphidium* in particular on the evolution of platyfishes through hybridization.

### Genome stabilization after hybridization

Homozygous regions in hybrid genomes evolved through recombination events in successive generations, and were finally fixed to the one or the other side of the parental ancestry, a process referred to as “genome stabilization”^74^. We refer to the ancestry from the “minor parent” as hybridization ancestry or foreign ancestry.

We found hybridization ancestry dispersed in different genomic areas following recent (Fig. 3I-L) and ancient (Fig. 3B-H) hybridization events. Meanwhile we failed to detect common enriched functions for foreign PCGs that were fixed in different hybridization events. This may reflect different selective forces and adaptations during the stabilization of different hybrid genomes or distinct outcomes of genetic drift in these independent hybridization events.

Different from the other species with hybridization history, *X*. *signum* exhibits a divergent pattern: while prevalent across chromosomes in other hybrid genomes, admixed signals are only steeply clustered mainly on chromosomes 15 and 18 in *X. signum* (Fig. 3I). The retention of only a small fraction of hybridization ancestry may be due to a limited adaptive value of the minority parent genome or structural features, like inversions. From zoogeographical reasons a recent hybridization of *X. signum* with *X. mayae* as the causation of this pattern can be excluded. Small clusters of hybridization ancestries were also observed on chromosome 11 and 18 of *X*. *mixei* (Fig. 3J). These clusters may contain functionally linked loci whose separation will be purged by selection against incompatibility. Alternatively, recombination events merely failed to break up these regions.

Based on whole-genome analysis, we detect hybridization in the lineage leading to *X*. *clemenciae*, *X*. *monticolus* and *X*. *mixei* (Fig. 3B-D). As expected from a shared ancestral hybridization event, we find that patterns of local ancestry are highly concordant across the genomes of these three species. This pattern is also observed in hybridization events involving *X*. *xiphidium* (Fig. 3E-H). An ancient hybridization was noted in cichlid fish, however the offspring species possess different combinations of biparental ancestry. This has been explained as being due to the rapid radiation of cichlid fish by sorting various combinations of biparental ancestry into different species^75–77^. Less rapid radiation in Southern swordtails may explain the stabilization before speciation in our case.

Molecular evolution of hybridization-derived regions has been studied by comparing the orthologous regions between two parental species, hence reflecting their evolution before the hybridization event^15^. Resulting from the hybridizations in ancestral lineages, common hybridization-derived regions shared between species provide us an opportunity to investigate their molecular evolution after hybridization. We found a significantly lower divergence and slower nonsynonymous substitution rate in hybridization-derived regions (Supplementary Fig. S10CE). This is congruent with the hypothesis that genomic regions with lower substitution rates, particularly at nonsynonymous sites, might be less likely to be associated with negative epistasis and incompatibilities between species^78–80^. It is worth noting that an opposite association was found in cases of recent hybridizations that occurred less than a few hundred generations ago^66^. We also detected a slightly higher value of dN/dS for coding genes with hybrid ancestry (Supplementary Fig. S10F). This could be explained by less constraint on such regions.

Our whole genome analysis uncovered that hybridization has occurred frequently during the evolution of *Xiphophorus*. Genome stabilization is a hallmark of molecular evolution of such hybrid-derived genomes. Whether the conservation of ancestral regions is a cause or consequence of divergence and speciation requires more detailed studies on the lineage specific adaptations in the 26 currently known species.

## Materials and Methods

### Sampling and genome sequencing

High molecular weight DNA was prepared from pooled soft organs of single individuals by a phenol/chloroform extraction procedure^81^. All individuals used for whole genome sequencing were raised at the fish facilities of the Biocenter of the University of Würzburg and the Xiphophorus Genetic Stock Center at Texas State University following approved experimental protocols through an authorization (568/300-1870/13) of the Veterinary Office of the District Government of Lower Franconia, Germany, in accordance with the German Animal Protection Law (TierSchG) and with an approved Institutional Animal Care and Use Committee protocol (IACUC 7381). Texas State University has an Animal Welfare Assurance on file with the Office of Laboratory Animal Welfare, National Institute of Health (A4147).

For *P. lacandonae* and *X. maculatus* Bp, muscle samples were used to obtain high molecular weight gDNA using the QIAGEN MagAttract HMW DNA kit (QIAGEN, Germantown, MD, USA) according to the manufacturer’s protocol. The quality and quantity of the gDNA were analyzed using a 5400 Fragment analyzer (Agilent Technologies, CA, USA) and Qubit 2.0 Fluorometer (Invitrogen, Life Technologies, CA, USA). DNA libraries were generated using the 10x Genomics Chromium technology according to the manufacturer’s instructions. Gel Bead-In-Emulsions (GEMs) were created from a library of Genome Gel Beads combined with 1.5 ng of gDNA in a Master Mix and partitioning oil, using the 10x Genomics Chromium Controller instrument with a microfluidic Genome chip (PN-120257). The GEMs were then subjected to an isothermal incubation step. Bar-coded DNA fragments were extracted and underwent Illumina library construction, as detailed in the Chromium Genome Reagent Kits Version 2 User Guide (PN-120258). Library yield was measured through the Qubit dsDNA HS assay kit (Thermo Fisher Scientific, Waltham, MA, USA). Library fragment size and distribution were measured using an Agilent 2100 Bioanalyzer High Sensitivity DNA chip (Santa Clara, CA, USA). The DNA was sequenced on a NovaSeq with a 2×150 bp read metric.

For *P. lacandonae*, muscle from the same sample were used for constructing a Hi-C chromatin contact map to enable a chromosome-level assembly. Tissue fixation, chromatin isolation, and library construction for Hi-C analysis were performed according to the manufacturer’s instructions (Dovetail Genomics, Chicago, USA)^82^. After checking the insert size, concentration, and effective concentration of the constructed libraries, final libraries were sequenced using the Illumina Novaseq platform (San Diego, CA, USA) with a 150-bp paired-end strategy.

### Genome Assembly

For genomes sequenced as Illumina short reads, we assembled each genome in two parallels, and chose the result with higher completeness or continuity. First, contigs were assembled using SOAPdenovo (Version 2.04)^83^, scaffolded using SSPACE (version 3.0)^84^ and arranged into chromosomes using cross_genome^85^. In parallel, contigs were assembled and scaffolded using Platanus (version: 1.2.4)^86^, and then arranged into chromosomes using cross_genome.

SOAPdenovo assemblies scaffolds by running de Brujin graph to merge all read-subsequences of length “*k*-mer”. We tried different *k*-mers (from 75-mer to 105-mer), chose the result with the longest N50 scaffold length, and then filled the gaps using GapCloser (version 1.12)^86^. The gap-filled scaffolds were then broken into contigs again at the N-linked positions. We removed those contigs that aligned to the mitochondrial genome and then rescaffoled them using SSPACE with parameters: -x 0 -z 200 -g 2 -k 2 - n 10.

Given the highly conserved synteny among genomes of Cyprinodontiformes^51^, we decided to assemble the scaffolds into chromosomes based on cross-species synteny using cross_genome. Highly qualified genome of *X*. *couchianus*, *X*. *maculatus*, *X*. *hellerii* and *X*. *malinche* respectively were used as the reference for Northern Platies, Southern Platies, Southern Swordtails and Northern Swordtails.

Platanus works well for assembling divergent heterozygous regions. Contigs were first assembled using command “platanus assemble” with parameter “-t 21 -m 80”. Contigs that aligned to the mitochondrial genome were removed and the rest were scaffolded using the command “platanus scaffold”. We then filled the gaps in scaffolds using “platanus gap_close” and GapCloser, and arranged the scaffolds into chromosomes using cross_genome.

Genomes sequenced as 10X linked-reads were assembled using Supernova (version 2.1.1)^87^ with parameters: –maxreads 298666666 –localcores=8. The assembly was output in style “pseudohap2” where two ‘parallel’ pseudohaplotypes were created and placed in separate FASTA files. From the two consensus assembly results we chose the one with higher completeness or continuity.

For *P. lacandonae*, Hi-C reads were mapped to the draft 10X genome assembly using HiC-Pro (v. 2.8.0) with default parameters^88^.

The completeness of each assembly was estimated using BUSCO (version 2.0.1)^89^ by the completed partition of the actinopterygii_odb9 database which contains the conserved gene set across actinopterygii (n=4584). The completeness could also be evaluated by comparison between estimated genome size and actual size of the sequences. The continuity of the assembly was evaluated by N50 scaffold length which was calculated by perl script assemblathon_stats.pl^90^.

### Genome Annotation

Genomes were annotated using a further developed and improved pipeline from our previous studies^91^. In brief, the assembly was masked at repeat regions, then mapped with protein sequences and RNA reads to collect homology and transcriptome gene evidence. Consensus gene models, given their high quality, were used to train the de novo gene predictor: AGUSTUS (version 3.2.3)^92^, which was then run with all collected gene evidence as hints to collect *ab initio* gene evidence. The final set of gene annotations was generated by synthesizing all three types of gene evidence.

In detail, to identify and mask repeats from the genome, we first screened the assembly for an automated discovery of transposable elements using RepeatModeler^48^. The consensus sequences of repeats were used as a *de novo* repeat library, together with Repbase and FishTEDB^93^. They were transferred to RepeatMasker for repeat identification and masking.

To collect homology gene evidence, we downloaded 455,817 protein sequences from the vertebrate database of Swiss-Prot (https://www.uniprot.org/statistics/Swiss-Prot), RefSeq database (proteins with ID starting with “NP” from “vertebrate_other”) and the NCBI genome annotation of human (GCF_000001405.39_GRCh38), zebrafish (GCF_000002035.6), platyfish (GCF_002775205.1), medaka (GCF_002234675.1), mummichog (GCF_011125445.2), turquoise killifish (GCF_001465895.1) and guppy (GCF_000633615.1). These protein sequences were aligned to the assembly using Exonerate (https://www.ebi.ac.uk/about/vertebrate-genomics/software/exonerate) and Genewise^94^, respectively, to predict gene location and intron/exon structures. To speed up Genewise, GenblastA was used a priori to find the rough location of the alignment on the assembly^95^.

To collect transcriptome gene evidence, RNA-seq reads from multiple tissues were cleaned using fastp^96^, and were mapped onto the assembly using HISAT^97^. The resulting bam file was then interpreted by StringTie for gene locations and structures^98^. In another parallel, transcript sequences were assembled based on the bam file using Trinity^99^. We then aligned these transcript sequences to the assembly for gene prediction using splign^100^.

AUGUSTUS was used for collecting de novo gene evidence. We first trained AUGUSTUS using BUSCO with the parameter “-long”^89^. In addition, those genes that were predicted repeatedly by Exonerate, Genewise, StringTie and splign were considered as being of high quality and were used to train AUGUSTUS for the second round. The trained AUGUSTUS was then run on the assembly with all homologous and transcriptome gene evidence as hints for an ab-initio gene prediction.

### Identification of orthologous genes

Orthologous genes were identified by sequence-similarity clustering followed by gene tree reconstruction within each cluster. Protein sequences of the longest transcript of each gene were pooled together and passed through an all-versus-all BLAST. Then for each two genes, an H-score was calculated from the BLAST score to index the sequence similarity^101,102^, so the genes could be clustered using Hcluster_sg with *P*. *lacandonae* as the outgroup^103^. Within each cluster a gene tree was reconstructed using TreeBeST v.0.5^104^. Based on which, the orthologous genes between species were typed as “*n* to *m*” orthology (*n* and *m* are positive integers; there are cases where *n*=*m*).

### Size dynamic of gene families

Gene families were identified by clustering genes with similar protein sequence (see Method section: identification of orthologous genes). For each gene family, we counted its member numbers in each species. Then CAFE5 was used to retrieve gene families that had been significantly expanded or contracted during the lineage evolution^105^.

### Mitochondrial genome tree

The mitochondrial genomes were assembled from the primary reads using MITObim and norgal^106,107^. MITObim assembles the genomes by baiting the reads and iteratively mapping them on a reference mitochondrial genome. The mitochondrial genomes of *X*. *couchianus*, *X*. *maculatus*, *X*. *hellerii* and *X*. *malinche* were downloaded from NCBI and used as the reference for Northern Platies, Southern Platies, Southern Swordtails and Northern Swordtails. For comparison and complementation, we also used norgal for de novo assemblies. As input for both assemblers, 10% clean reads were sampled from the whole-genome sequencing (WGS) using BBmap^108^.

Mitochondrial genomes were annotated using the MitoAnnotator pipeline of the Mitofish database^109^. DNA sequence alignment of each protein coding gene, rRNA and tRNA was made using MACSE for coding sequences and MUSCLE for non-coding sequences^110,111^. We manually screened the alignments to check for potential assembly or annotation errors, which were then curated by selecting a better assembly from MITObim and norgal result. With respect to annotation errors in protein coding genes, we selected the better assembly, and retrieved the coding sequence by aligning homologous protein sequence on the DNA sequence using Genewise. After curation, alignment gaps were removed using Gblocks^112^.

The phylogenetic tree based on mitochondrial genomes was built using RAxML and MrBayes respectively^113,114^. RAxML infers phylogeny relationships under maximum likelihood. As input, we concatenated the DNA alignments of the 13 protein coding genes, 2 rRNAs and 22 tRNAs into a single giant alignment. In addition, the region information of each gene was passed to RAxML in a partition file for the partitioned analysis. This resulted in an 15,515 bp long alignment with partitions as follow: ATPase-6 (1-681), ATPase-8 (682-846), COIII (847-1629), COII (1630-2319), COI (2320-3876), Cyt-b (3877-5016), ND1 (5017-5976), ND2 (5977-7020), ND3 (7021-7365), ND4L (7366-7662), ND4 (7663-9042), ND5 (9043-10869), ND6 (10870-11382), 12S-rRNA (11383-12319), 16S-rRNA (12320-13969), tRNA-Ala (13970-14038), tRNA-Arg (14039-14107), tRNA-Asn (14108-14180), tRNA-Asp (14181-14251), tRNA-Cys (14252-14315), tRNA-Gln (14316-14386), tRNA-Glu (14387-14455), tRNA-Gly (14456-14526), tRNA-His (14527-14596), tRNA-Ile (14597-14663), tRNA-Leu2 (14664-14735), tRNA-Leu (14736-14809), tRNA-Lys (14810-14882), tRNA-Met (14883-14950), tRNA-Phe (14951-15018), tRNA-Pro (15019-15087), tRNA-Ser2 (15088-15156), tRNA-Ser (15157-15227), tRNA-Thr (15228-15300), tRNA-Trp (15301-15373), tRNA-Tyr (15374-15443), tRNA-Val (15444-15515). The analysis was performed under the General Time Reversible (GTR) + Gamma phylogenetic model with 1,000 rapid bootstrap tests.

MrBayes infers phylogeny using Bayesian methods. The concatenated alignment was used also as the input data. We ran the inference under a General Time Reversible model with a proportion of invariable sites and a gamma-shaped distribution of rates across sites by setting parameter “lset nst=6 rates=invgamma”. Three runs starting from a different tree were launched with six chains for each running 50 million generations. The results were sampled each 1000 generations and the first 25% samples were discarded as burn-in.

### Phylogenomic tree based on whole genome alignment (WGA)

The multiple WGA across all species was built by merging pairwise genome alignments of each species to *X*. *maculatus*. We used minimap2 for the pairwise genome alignments with parameter “-cx asm20 –cs=long”^115^. The alignments were then refined using Genome Alignment Tools from Hiller lab as follows^116^: alignments were first chained up using axtChain, the unaligned loci flanked by aligning blocks were then re-aligned using patchChain.perl, newly-detected repeat-overlapping alignments were incorporated into the alignment chains using RepeatFiller. To improve the specificity of alignments, the obscure local alignments were detected and removed using chainCleaner. We then used chainNet to collect alignment chains hierarchically to capture only orthologous alignments. Finally, pairwise genome alignments were merged into the multiple WGA using MULTIZ^117^. The final WGA contains 340,357 alignment blocks with a mean length of 1,214 bp, the longest of ∼34 kb and the sum of ∼413 Mb, spanning ∼58% of the genome.

To reconstruct a phylogenomic tree based on the WGA, we removed alignment blocks shorter than one kb to avoid potential false-positive-alignments. The remaining alignment blocks were then trimmed using trimAl and then concatenated into a ∼342 Mb long alignment^118^. The alignment was passed to RAxML for a maximum likelihood inference of the phylogenomic tree under GTR+Gamma model with 1,000 rapid bootstraps. The bootstrap values of nodes were all at 100%.

To construct a coalescent tree, we sampled 19,111 alignment blocks that are at least 1 kb long and 20 kb away from each other. From each of the blocks we randomly cut out a 1kb window to infer the maximum likelihood tree using RAxML under GTR+Gamma model with 100 rapid bootstraps. Trees with none of the nodes supported over 75% were removed, nodes with bootstrap value <75% were collapsed. The final coalescent tree was built based on the filter trees using ASTRAL-III^119^.

### Phylogenomic tree based on protein coding genes

3259 one-to-one orthologous genes were identified across all species based on sequence similarity followed by synteny confirmation. With these orthologies, we reconstructed the phylogenomic tree using RAxML with the concatenated protein sequences, coding sequences and 4DTV (fourfold degenerate sites) sequences respectively. Evolutionary model was automatically searched for protein sequences with parameter “-m PROTGAMMAAUTO”, while set to GTR+Gamma for coding and 4DTV sequences. Node confidence of each tree was accessed by 200 bootstraps.

### Inference of hybridization history

The whole genome alignments of four target species were constructed using minimap2 and MULTIZ, then a 1kb-long window was extracted from each of the alignment blocks that are at least 1 kb long and 20 kb away from each other. For each window, we applied AU-test to select the optimal tree topology from the three candidates. The site-wise likelihoods were first calculated using RAxML and then the AU-tests implemented in Consel 0.20^120^. At last, trees with AU *P*-value higher than 0.95 were included for the count and comparison.

We also applied Patterson’s D-statistic on these alignment windows to detect the gene flow between two species. Patterson’s D-statistic is also known as ABBA-BABA statistic, where alignment sites ABBA refers to those at which species 2 and 3 share a derived allele “B”, while species 1 has the ancestral state “A”, as defined by the outgroup: species 4. Likewise, BABA refers to those sites at which species 1 and 3 share the derived state. The counts of the two type sites would equal each other when there is no gene flow. This null hypothesis was tested using a two sample z-test. Standard error (SE) of value D was determined using jack-knife resampling method^121^.

AU-test and the D-statistic can infer the hybridization history when only three species were targeted. To calculate the D and f4-ratio statistics across all combinations of the studied species, we also implemented Dsuite^56^. At last, specific introgressed loci were retrieved with parameter “Dinvestigate -w 50,5” using fdM index, which may flaws in simulations but it is the best option still for our analyses.

### Retrieve sequences for *xmrk* and *egfrb*

First, we retrieved the sequences of *xmrk* and *egfrb* for each species from the genome assembly through the result of genome annotation and *ab inito* homology search. Then for those species with no *xmrk* found in the assembly, we made an effort to recover it directly from the reads. Protein sequences of *xmrk* from *X*. *maculatus* (XP_023181635.1 and XP_023185812.1) were used as the reference. Reads were aligned to the reference using diamond^122^. We then assembled the retrieved reads using cap3 and then translated the resulting contigs into protein sequences using GeneWise^94,123^. At last, for each exon of a reference, the contig match the best were kept and assembled into the final sequence.

## Data availability

Raw reads of the whole genome sequencing in the study are available in SRA under accession number PRJNA972672. Assemblies and annotations are available in figshare under DOI 10.6084/m9.figshare.23596515.

## Author contribution

MaS and AM designed the study and wrote the paper. KD performed the assemblies, annotations, all bioinformatic analyses and drafted the manuscript. HP sequenced and assembled the genome of *P. lacandonae* and *X. maculatus* Bp. RBW, WCW, MoS, YL and AM provided raw data and assembled genomes. All authors interpreted the results and participated in manuscript writing.

## Acknowledgements

We thank the *Xiphophorus* Genetic Stock Center for providing several fish samples used in this study. This work was supported by NIH R24OD031467, NIH R15CA223964, NIH 1R35GM133774 and CPRIT RP200657. Computational work associated with the study was performed on the Learning, Exploration, Analysis, and Processing (LEAP) next-generation High-Performance computing cluster at the Texas State University, San Marcos, TX. This work was supported by an ERC advanced grant (293700) to AM.

