## Supplementary Figures for "Phylogenomics analyses of all species of Swordtails (Genus *Xiphophorus*) highlights hybridization precedes speciation"

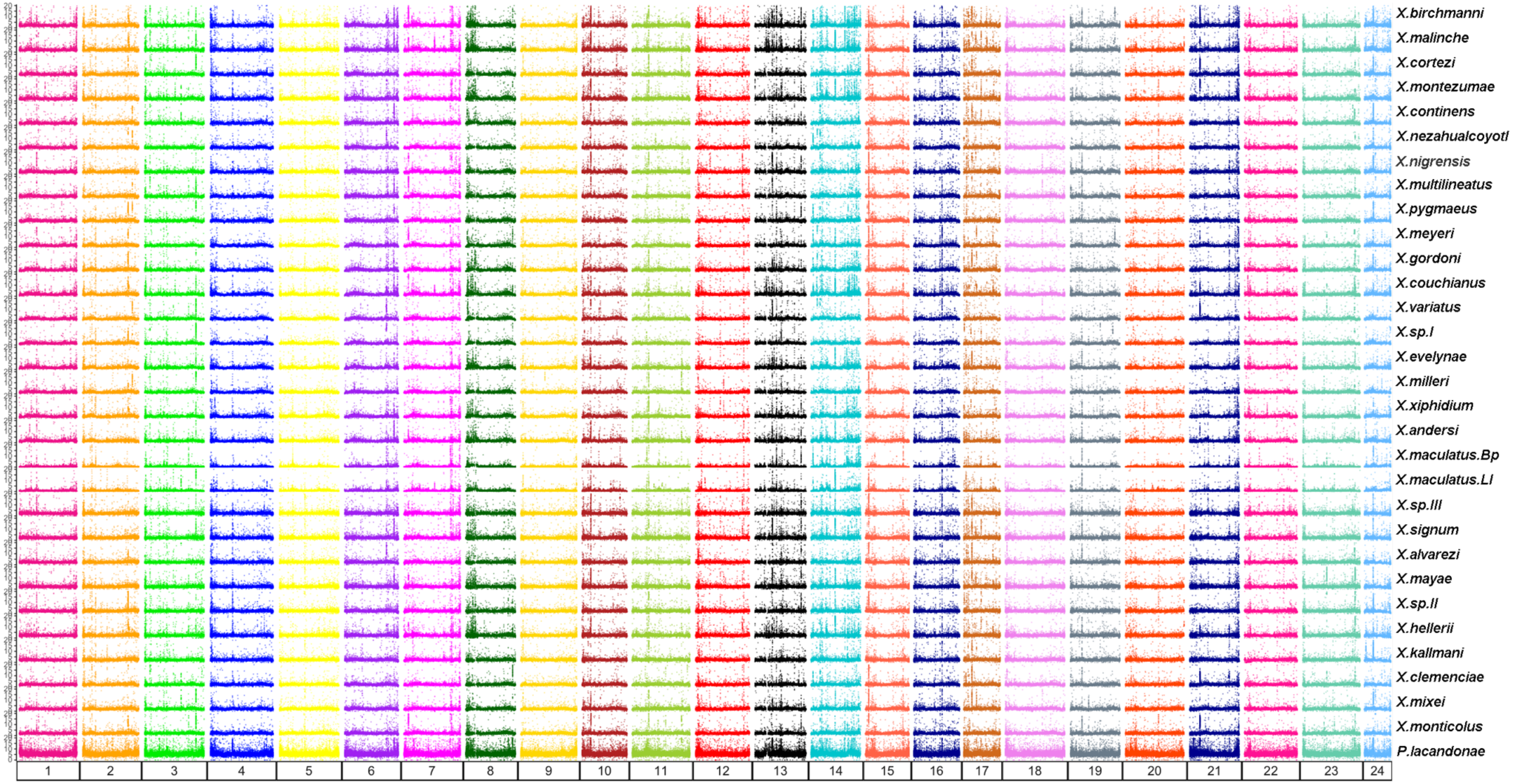


**Supplementary figure S1. Plots revealing the sequence difference between *X. maculatus* and all other species across chromosomes.** Sequence difference was calculated in 10kb sliding windows as the percentage of SNP and indels.

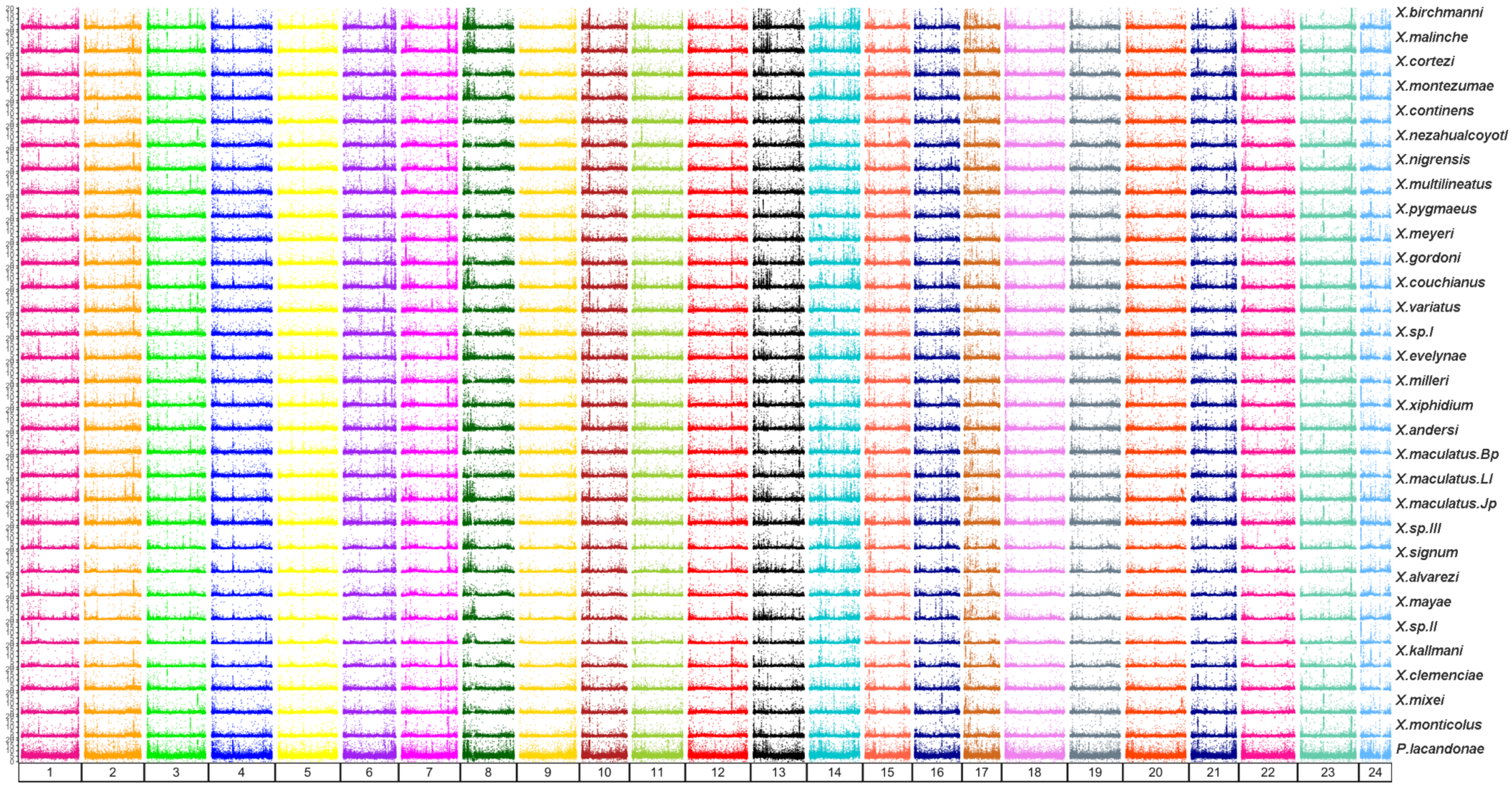


**Supplementary figure S2. Plots revealing the sequence difference between *X. hellerii* and all other species across chromosomes.** Sequence difference was calculated in 10kb sliding windows as the percentage of SNP and indels.

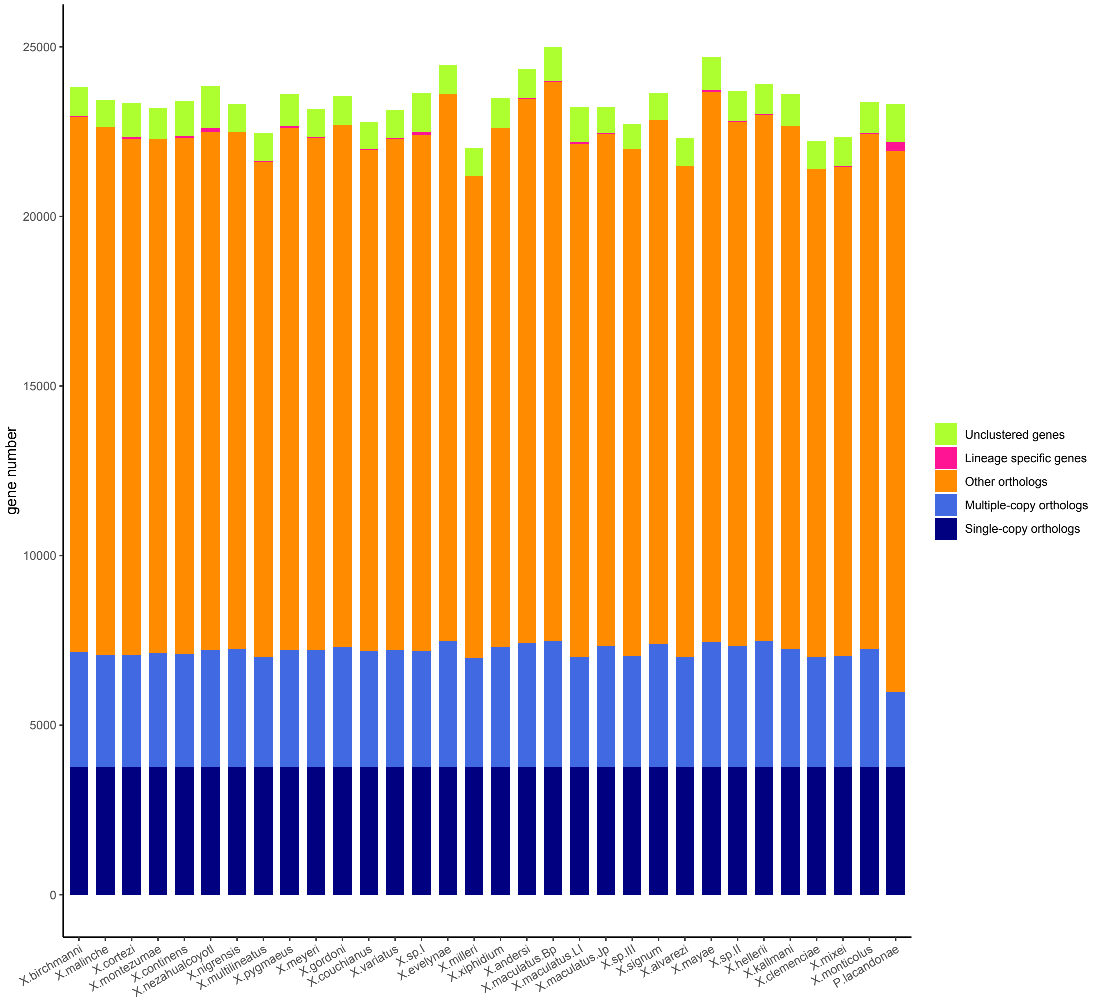


**Supplementary figure S3. Bar plots revealing the content of orthologous protein coding genes across species.**

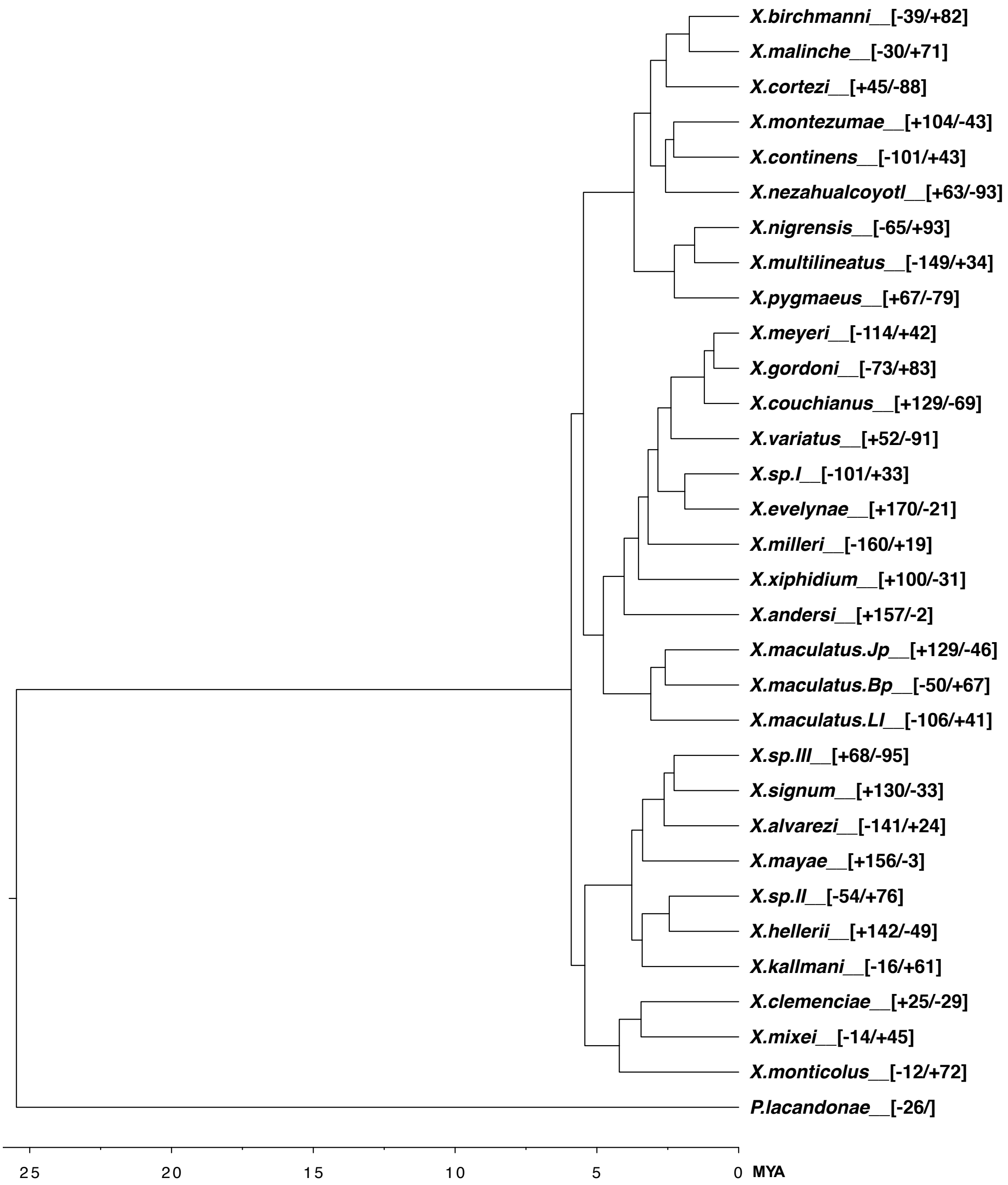


**Supplementary figure S4. Phylogeny tree revealing the time of branch divergence and gene family expansion and contraction on each lineage.** Number after + is the number of significantly expanded gene family; after -, the significantly contracted.

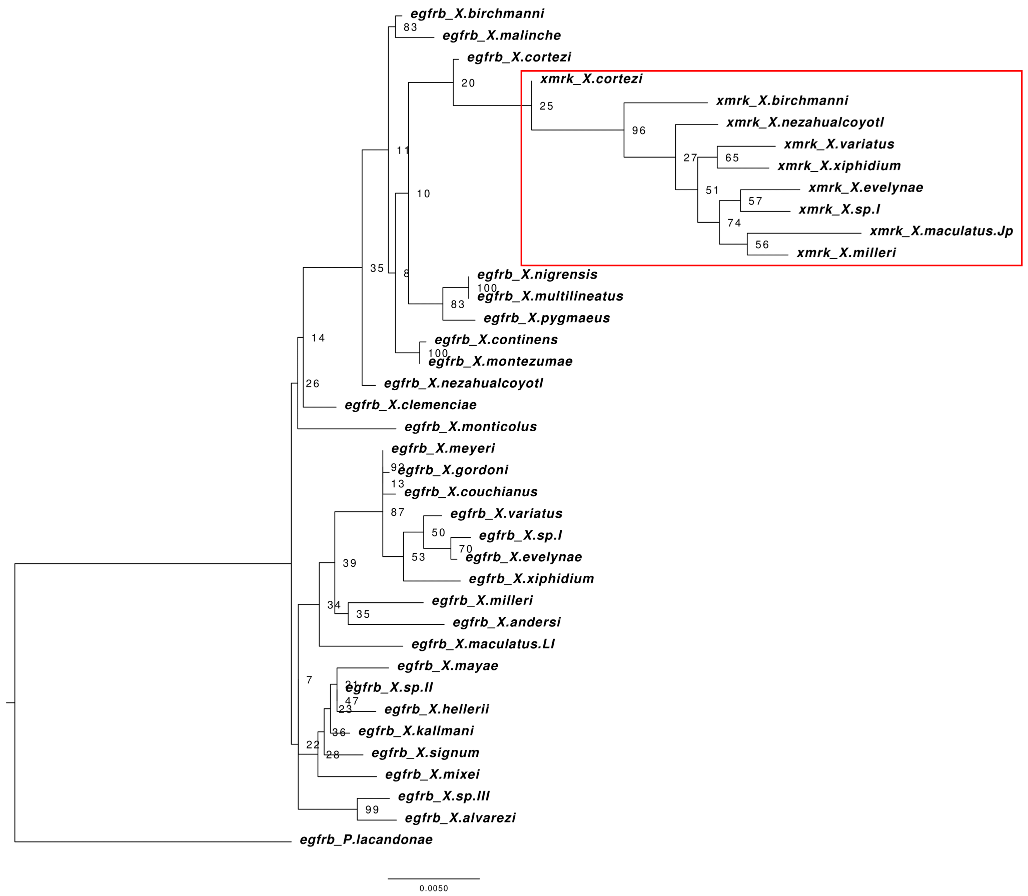


**Supplementary figure S6. Phylogeny tree of *xmrk* constructed using coding sequence with maximum-likelihood method. Numbers are bootstrap support value.**

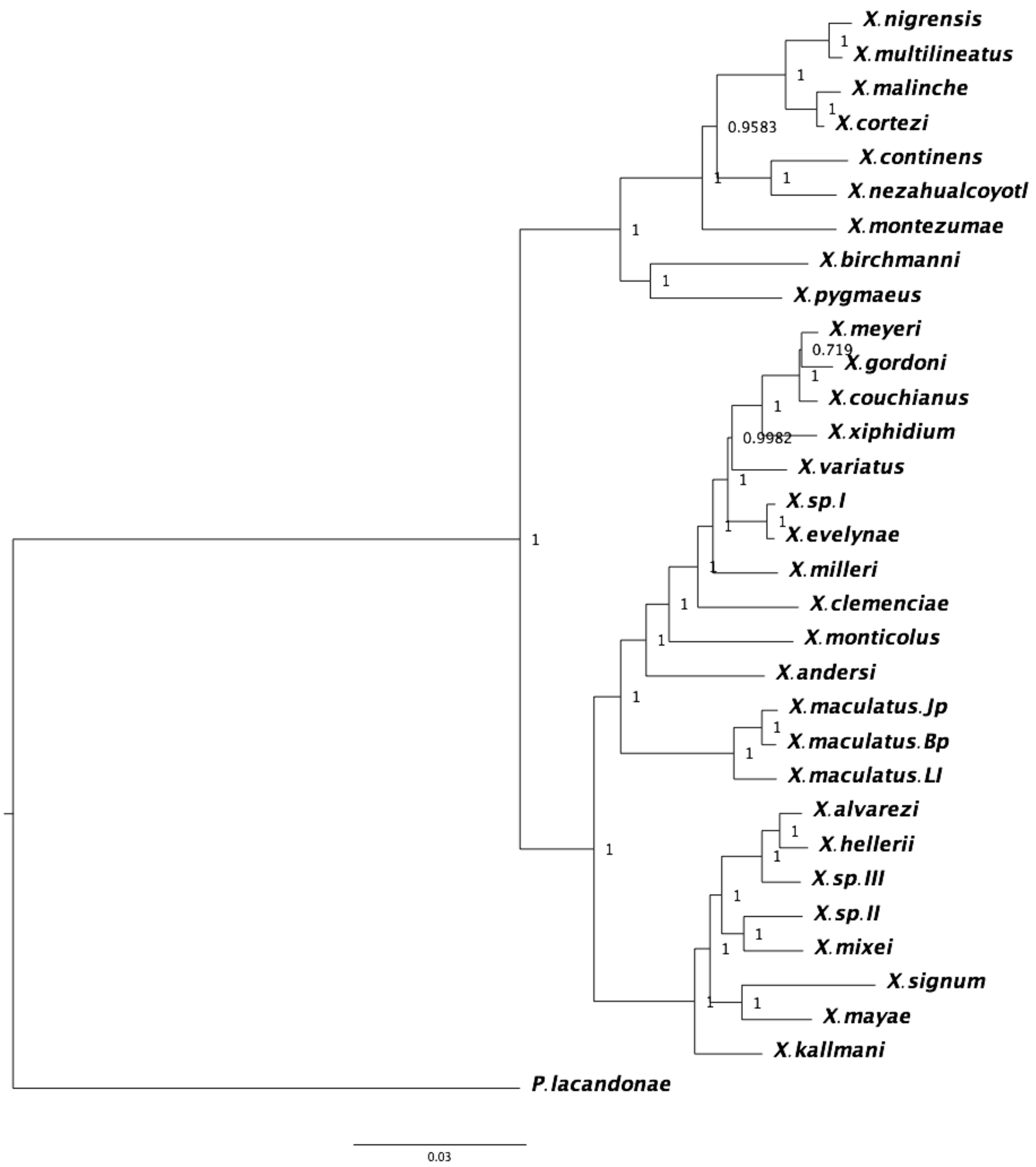


**Supplementary figure S7. Mitochondrial phylogeny tree constructed using mitochondrial sequences with the method of bayesian inference.**

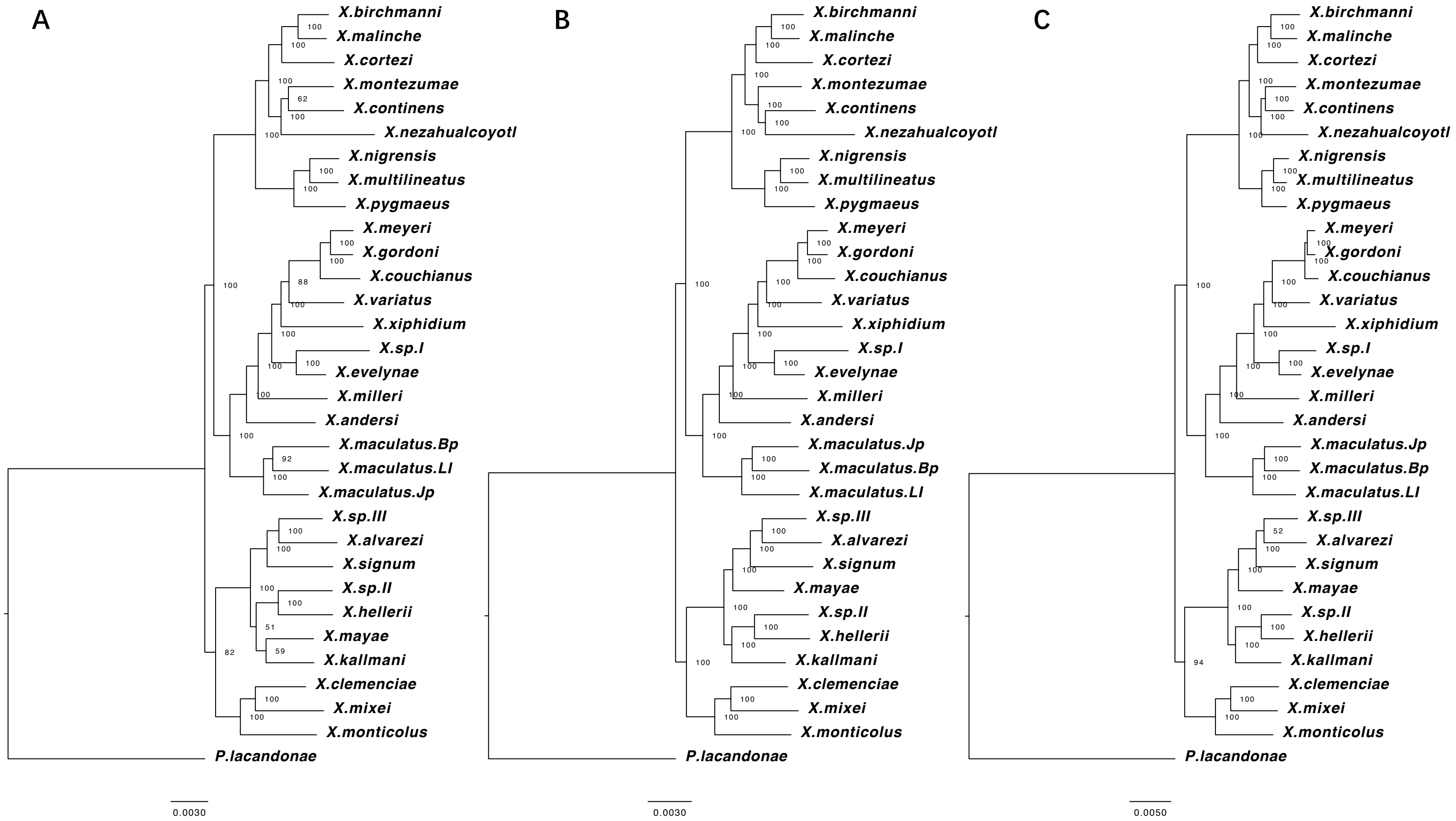


**Supplementary figure S8. Phylogenetic tree constructed using max-likelihood method based on protein sequences (A), coding sequences (B) and 4DTV sites (C) of 3,259 one to one orthologous genes.** Number on the nodes represents bootstrap support value.

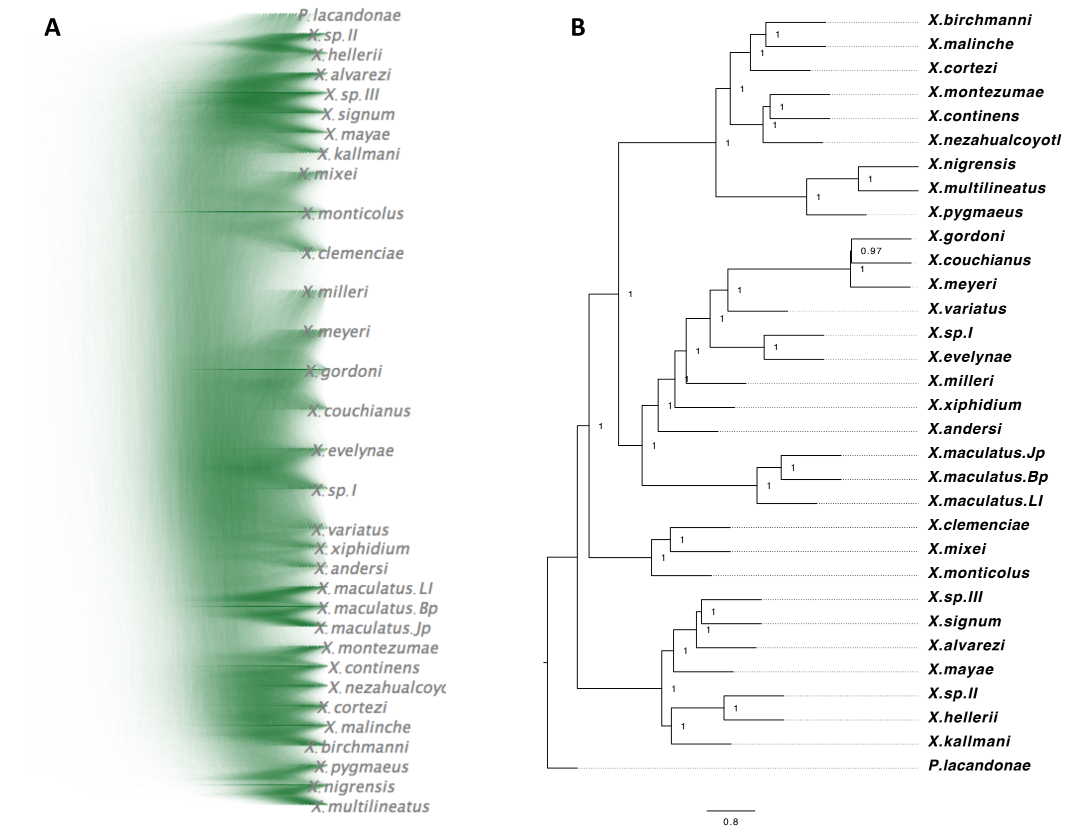


**Supplementary figure S9. Coalescent phylogeny.** (A) Density tree revealing ~12k trees made from sequences of genomic loci. (B) Phylogenetic trees constructed using a coalescent method.

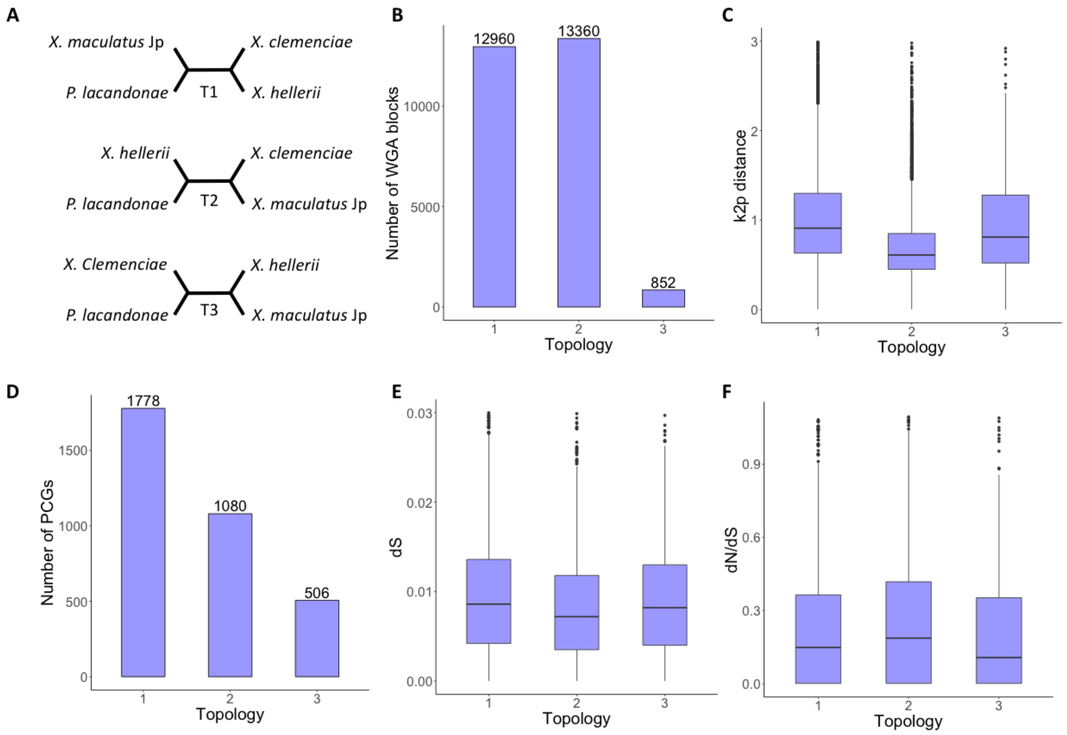


**Supplementary figure S10. Incongruence phylogenies and evolutionary character of admixed genomic loci for hybridization between *X. clemenciae* and *X. maculatus*.** (A) Phylogenetic topologies with T1 as the genome-wide consensus tree, T2 suggesting ILS or hybridization between *X. clemenciae* and *X. maculatus* and T3 suggesting ILS or hybridization between *X. hellerii* and *X. maculatus*. (B) Number of WGA regions past AU test yielding different topologies. (C) k2p distance between *X. clemenciae* and *X. monticolus* for WGA regions corresponding to different-topologies. (C) Number of PCGs past AU test yielding different topologie. (E) dS values of PCGs corresponding to different-topologies. (F) dN/dS values of PCGs corresponding to different-topologies.

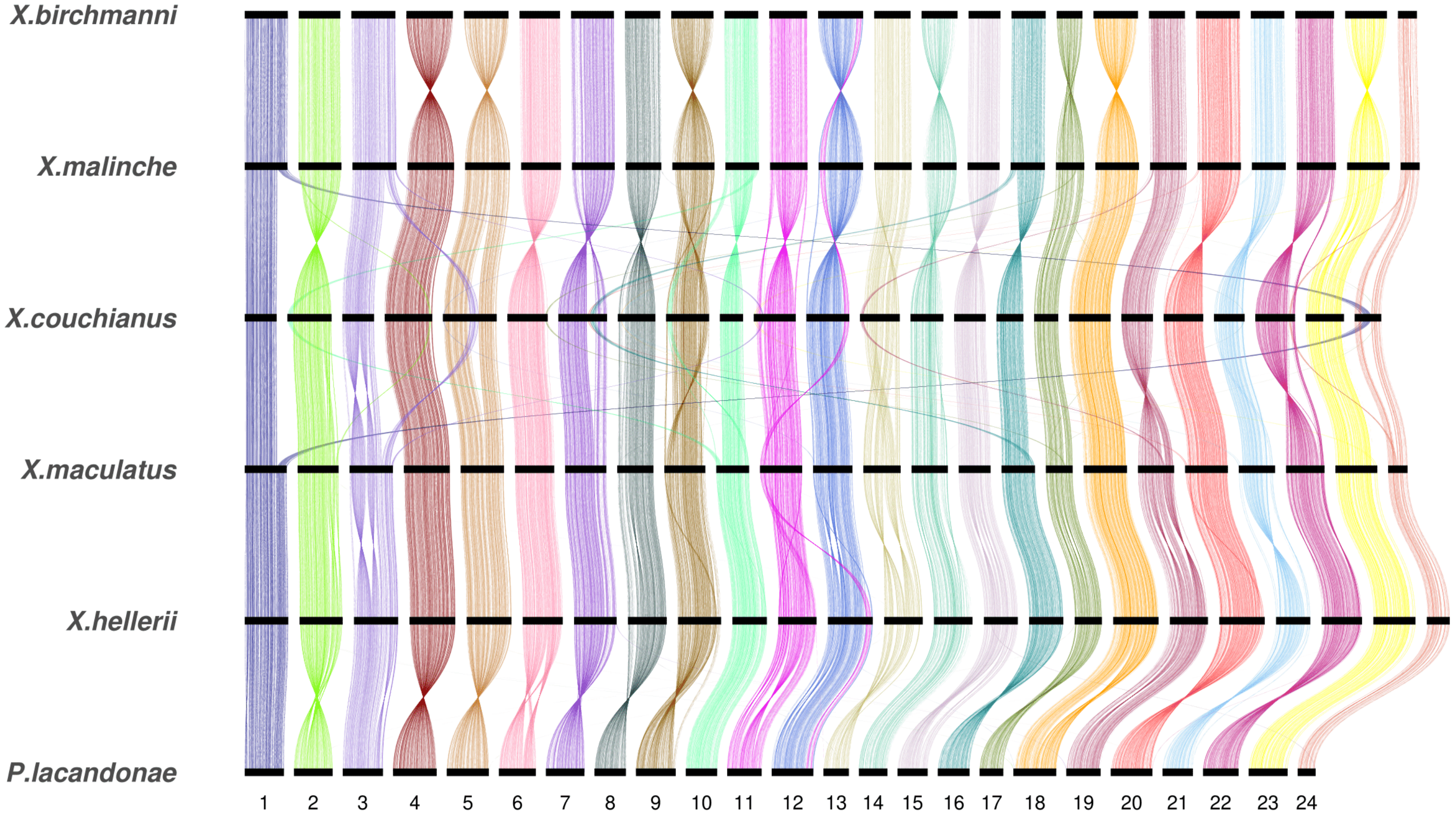


**Supplementary figure S5. Sankey plots showing conserved synteny across the six species with chromosome-level assemblies.** Black bars represent the chromosome and threads link the one-to-one orthologous genes across species.

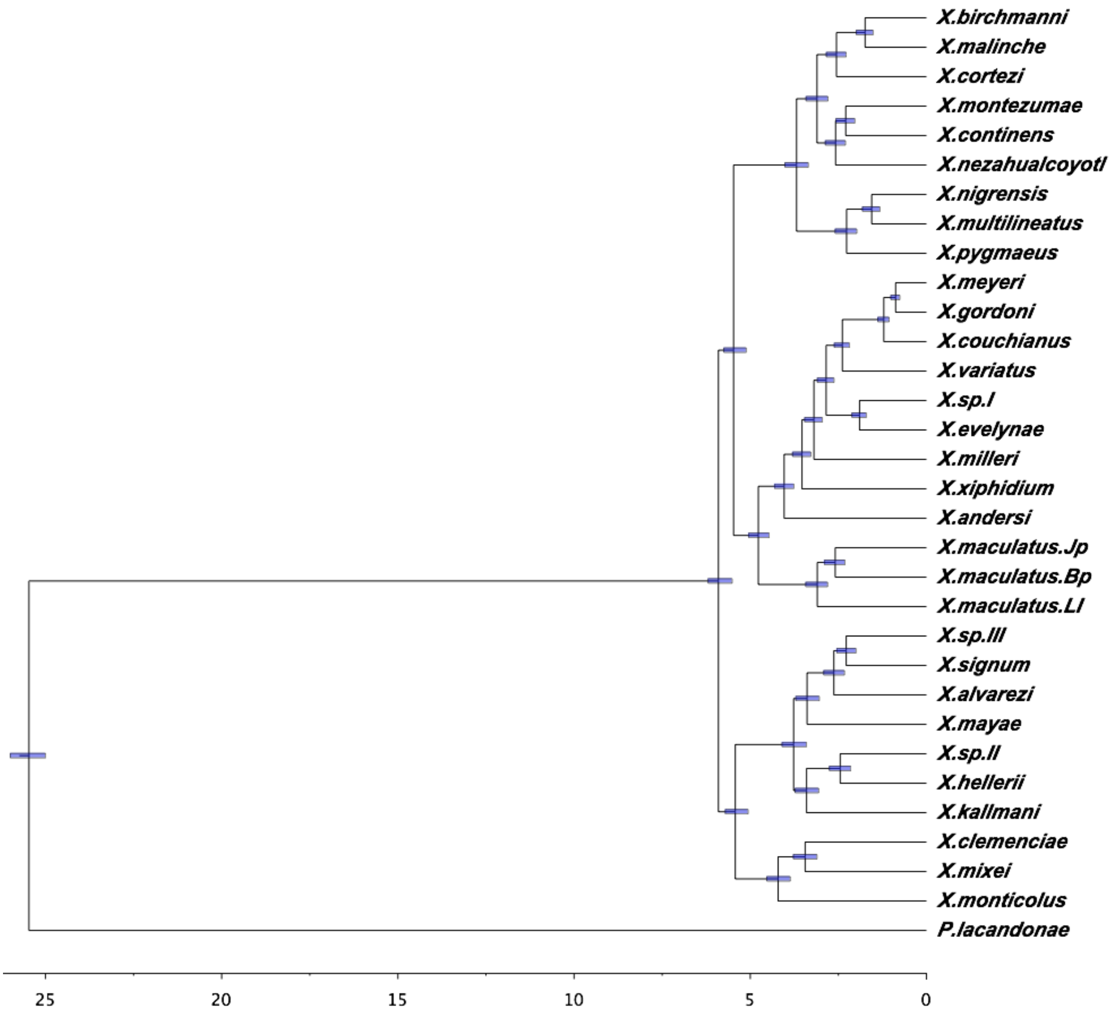


**Supplementary figure S11. Phylogeny tree showing the time schema for the evolution of *Xiphophorus*.** Unit on X axis is million years ago (MYA). Blue bar on each node represents the 95% confidence interval (CI) for the time estimation.
